# Parallel EEG assessment of different sound predictability levels in tinnitus

**DOI:** 10.1101/2023.07.25.550472

**Authors:** Pia Brinkmann, Jana V. P. Devos, Jelle H. M. van der Eerden, Jasper V. Smit, Marcus L. F. Janssen, Sonja A. Kotz, Michael Schwartze

## Abstract

**Objective:** Tinnitus denotes perception of a non-environmental sound and might result from aberrant auditory prediction. Successful prediction of formal (e.g. type) and temporal sound characteristics facilitates the filtering of irrelevant information (“sensory gating”, SG). Here, we explored if and how parallel manipulations of formal and temporal predictability affect sensory gating in persons with and without tinnitus.

**Methods:** Age-, education- and sex-matched persons with and without tinnitus (N = 52) participated and listened to paired-tone “oddball” sequences, varying in formal (standard vs. deviant pitch) and temporal predictability (isochronous vs. random timing). EEG was recorded from 128 channels and data were analyzed by means of temporal spatial principal component analysis (tsPCA).

**Results:** SG was observed in P50- and N100-like activity (amplitude suppression for the 2^nd^ tone in the pair) in both timing conditions and groups. Correspondingly, deviants elicited overall larger amplitudes than standards. However, only in persons without tinnitus N100-like activity in response to deviants was enhanced with isochronous relative to random timing.

**Conclusions:** Persons with tinnitus do not benefit similarly as persons without tinnitus from temporally predictable context in deviance processing.

**Significance:** The current results indicate altered temporal sensitivity and selective attention allocation in persons with tinnitus.

**Highlights:**

- Persons with tinnitus display altered auditory predictions affecting the processing of unexpected auditory input
- Position predictions did not differ between persons with tinnitus and without
- Temporal predictability facilitated deviance processing for P50-like activity in persons with tinnitus and without

## 1. Introduction

Predicting the type and timing of upcoming events guides goal-directed behavior and is key to efficiently adapting to changes in an abundant sensory environment. In audition, predictions can take different forms. They are either based on formal (type) or temporal (timing) stimulus characteristics (Bendixen, SanMiguel, & Schröger, 2012; Mauk & Buonomano, 2004; Tavano, Widmann, Bendixen, Trujillo-Barreto, & Schröger, 2014). Formal predictions refer to the spectral information that is conveyed in an acoustic stimulus (Schwartze, Tavano, Schröger, & Kotz, 2012), while temporal predictions refer to the points in time when a stimulus event occurs (Schwartze et al., 2012). Nested in this basic distinction, predictions can be based on specific stimulus arrangements such as the repetitive binary stimulus grouping that is commonly used in ‘sensory gating’ (SG) studies to induce position predictions.

The fundamental importance of predictions is particularly evident when their underlying mechanisms change in pathological conditions. One condition in which altered auditory predictions seem to play a role is tinnitus (Brinkmann, Kotz, Smit, Janssen, & Schwartze, 2021; De Ridder et al., 2014; Hullfish, Sedley, & Vanneste, 2019; Roberts, Husain, & Eggermont, 2013; Sedley, Friston, Gander, Kumar, & Griffiths, 2016). Tinnitus is typically described as the ‘ringing in the ears’ and often experienced as a constant sound in the absence of any physical sound source (Axelsson & Ringdahl, 1989; Roberts et al., 2010). The most prominent risk factors for developing tinnitus are aging and hearing loss (Roberts et al., 2010). In the general population, its prevalence ranges from 10-14% in middle-aged adults and further increases with age (Jarach et al., 2022; Langguth, Kreuzer, Kleinjung, & De Ridder, 2013). Persons with tinnitus are either characterized by decompensated or compensated tinnitus. The first group suffers from tinnitus while the second is not substantially affected by it.

Taking a predictive coding perspective, chronic tinnitus might display altered default predictions, meaning that in chronic tinnitus, default predictions in audition change to represent ‘something’ instead of ‘silence’ (Hullfish et al., 2019; Sedley, Alter, Gander, Berger, & Griffiths, 2019). Alternatively, peripheral or subcortical tinnitus models propose that tinnitus might stem from aberrant cochlear activity (Mulders & Robertson, 2009) or impaired noise-cancellation due to malfunctioning in limbic structures (Rauschecker, Leaver, & Mühlau, 2010). It was suggested that tinnitus results from discrepant expected and actual auditory input in interaction with attention (Roberts et al., 2013). Although the exact interplay of these factors is unknown, discrepancy of this kind can lead to auditory phantom perception. As tinnitus is typically perceived as a tone with a constant pitch, it is assumed that formal predictions are most affected (Sedley et al., 2019). Additionally, altered temporal predictions in tinnitus have first been suggested in thalamocortical dysrhythmia (De Ridder, Vanneste, Langguth, & Llinas, 2015) and were subsequently discussed in a predictive network hypothesis (Brinkmann et al., 2021). Accordingly, distinguishing dimensions of auditory prediction (i.e., formal-, temporal-, and position-predictions) combined with a differential assessment of their function in tinnitus might lead to a better and more comprehensive understanding of tinnitus beyond the level of formal predictions.

The high temporal resolution of the electroencephalogram (EEG) provides an excellent tool for investigating auditory predictions. The P50 and N100 event-related potential components (ERPs) that peak around 50 and 100 ms post-stimulus respectively are responsive to prediction. SG is often described as a predictive filtering mechanism and investigated by presenting pairs of identical sound stimuli. The response to the second stimulus leads to a suppressed P50 ERP response (i.e., ‘gating out’), and indicates that the first stimulus is predictive of the second one (Adler et al., 1982; Cromwell, Mears, Wan, & Boutros, 2008).

The P50 is generated in temporal and frontal cortices and interpreted as an indicator of SG (Korzyukov et al., 2007; Smith, Boutors, & Schwarzkopf, 1994). P50 SG is conceived as a pre-attentional mechanism and mainly reflects sensory processes (Jerger, Biggins, & Fein, 1992; Kho et al., 2003). Modulation of the P50 response in SG might thus indicate the relative success of filtering out goal-irrelevant information (Jones, Hills, Dick, Jones, & Bright, 2016). Accordingly, stronger P50 suppression is associated with better attentional orienting and inhibition (Wan, Friedman, Boutros, & Crawford, 2008).

The N100 is generated in the supratemporal plane of the auditory cortex (Näätänen & Picton, 1987) but also in the frontal cortex (Giard et al., 1994). The N100 reliably indicates formal predictions as assessed in pitch-based “oddball” paradigms (Segalowitz & Barnes, 1993) and stands more for (selective) attentional processes (Thornton, Harmer, & Lavoie, 2007). Previous N100 research, that manipulated formal and temporal stimulus predictability, found differences between temporal and formal conditions (Schwartze, Farrugia, & Kotz, 2013). The N100 response to predictable stimuli becomes smaller over time and the interval between the stimuli is a factor that determines this decrease (Budd, Barry, Gordon, Rennie, & Michie, 1998). Taken together, the existing evidence suggests that the P50 and N100 might indicate different stages of predictive sensory filtering, with successful SG leading to better task performance and protected higher-order cognitive functioning (Lijffijt et al., 2009; Venables, 1964).

Neurophysiological SG studies in adults with tinnitus have produced inconclusive results (Campbell, Bean, & LaBrec, 2018; Dornhoffer, Danner, Mennemeier, Blake, & Garcia-Rill, 2006). For example, it has been shown that the SG difference index of the Pa component that precedes the P50, correlates negatively with tinnitus severity, i.e., more severe tinnitus reduced the Pa suppression in response to the second tone in a tone pair (Campbell et al., 2018). However, no significant differences were observed between persons with tinnitus and those without for Pa, P50, N100 or P200 gating effects (Campbell et al., 2018). Notably, participants in this study only experienced mild tinnitus symptoms as assessed by the tinnitus handicap inventory (THI) (Newman, Jacobson, & Spitzer, 1996). Another study investigated P50 suppression in persons with and without tinnitus and similarly did not report group differences (Dornhoffer et al., 2006). Habituation to repetitive auditory input seems to be reduced in persons with decompensated tinnitus as evident in N100 and P200 amplitude differences (Walpurger, Hebing-Lennartz, Denecke, & Pietrowsky, 2003). Sedley et al. (2019) manipulated formal predictability in a roving standard oddball paradigm and observed no differences between persons with or without tinnitus comparing their response to standard and deviant tones in the P50, while deviants evoked larger N100 responses in both groups. Thus, previous evidence shows reduced N100 - P200 amplitude differences during continuous repetitive stimulus presentation in persons with decompensated tinnitus (Walpurger et al., 2003), that increased tinnitus burden might be linked to impaired Pa suppression (Campbell et al., 2018), while for P50 SG no group differences were found (Dornhoffer et al., 2006). Research focusing on P50 and N100 responses as indices for stimulus type or temporal predictability in tinnitus thus likely reflect the heterogeneity of the condition (Cederroth et al., 2019).

Tinnitus has previously been investigated by means of stimulus sequences that incorporated manipulations of either formal or position prediction in isolation (Campbell et al., 2018; Dornhoffer et al., 2006; Sedley et al., 2019). However, experimental paradigms that allow the parallel assessment of different stimulus type dimensions and timing are needed to obtain a better understanding of how prediction impacts tinnitus heterogeneity. Such a comprehensive approach could allow differentiating individual prediction capacities and identify which dimensions are dysfunctional in tinnitus. As SG is a filtering mechanism that might operate on all predictability dimensions, it also allows exploring possible interactions between temporal-, formal- and position-based SG. Finally, linking this approach to specific ERP markers, such as P50 and N100, might allow decomposing the underlying mechanisms of selective attention and inform how they look in persons suffering from tinnitus.

Along these lines, the current study assessed if and how ERP markers of auditory predictions are altered in persons with and without tinnitus. The experimental setup incorporated a paired-tone oddball design that simultaneously manipulated formal and temporal stimulus features to differentiate and relate different aspects of predictability. This setup combined elements of previous studies (Schwartze et al., 2013; Schwartze, Rothermich, Schmidt-Kassow, & Kotz, 2011) with the aim to verify if previous findings could be reproduced. It was expected that tinnitus alters SG efficiency. Accordingly, it was hypothesized that successful SG for position, deviance, and regularity dimensions would result in smaller P50 and N100 amplitudes for predictable stimuli but that dysfunctional SG in tinnitus would lead to increased P50 and N100 amplitudes.

## 2. Methods

The study was approved by the ethics committee of the Maastricht University Medical Center + (MUMC+) with the code 2019-0970. Due to the COVID pandemic, data collection was paused and then continued intermittently under strict safety regulations.

### 2.1 Recruitment and inclusion

Participants were recruited via leaflets, word of mouth, and an existing database of persons with tinnitus. Persons with and without (subjective) tinnitus were included when they were between 18 and 69 years old and had an audiogram and a bilateral high tone Fletcher Index lower than 60 dB. Exclusion criteria were objective tinnitus (i.e., pulsatile tinnitus), a maximum air-bone gap of more than 20 dB, or a history of ear surgery, brain surgery or brain/ear implants. If available, participants provided their audiograms, all but seven obtained within the last year, otherwise an audiogram was obtained by trained personnel before testing. The two groups were matched for sex at the participant level and for age and education at the group level.

### 2.2 Participants

Fifty-two persons with tinnitus and without participated (**Table 1**.). Education was scored on 8 levels in persons with and without tinnitus, where 8 was the highest level. Handedness was assessed with the Dutch handedness questionnaire (van Strien, 2003) on a scale ranging from −10 (extreme left-handedness) to 10 (extreme right-handedness) for persons with tinnitus and without. Hearing loss (HL) was assessed using the average pure tone audiometry (PTA) in dB for the left and right ears for persons with tinnitus and without. Tinnitus burden was assessed with the Dutch version of the Tinnitus Questionnaire (TQ) (Goebel & Hiller, 1994; Meeus, Blaivie, & Van de Heyning, 2007) in the tinnitus group (*M_TQ_* = 37.3, *SD_TQ_* = 17.6), scores between 31 and 46 points indicate mild tinnitus burden (Grade II). Tinnitus duration was assessed in months (*M_TinDuration_* = 82.8, *SD_TinDuration_* = 76.4). The tinnitus group suffered from chronic tinnitus, considering that tinnitus is ‘chronic’ if it is experienced for at least three months (Snow, 2004).

**Table 1.** Demographics of the study participants.

| **Variables** | **with Tinnitus** | **without Tinnitus** |
| --- | --- | --- |
| Total Number (n) | 27 (7 female) | 25 (6 female) |
| Age ± SD | 50.6 ± 14.1 | 44.8 ± 16.3 |
| Education (0-8) | 4.5 ± 1.6 | 5.3 ± 1.7 |
| Handedness (-10 – 10) | 8.2 ± 3.7 | 4.6 ± 8.2 |
| Pure tone average (PTA) in dB for both ears | 23.2 ± 12.3 | 11.7 ± 12.9 ** |

\*\* $p \leq .01$

### 2.3 Materials

#### 2.3.1 Procedure

Upon arrival at the laboratory, participants signed the informed consent and performed the handedness questionnaire, answered questions about their tinnitus (i.e., tinnitus duration etc.), and filled in the TQ. They then entered an electronically shielded and soundproof booth for the EEG recordings.

#### 2.3.2 Experimental design and stimuli

The two stimulus sequences each consisted of 1152 standard (600 Hz) and 288 deviant (660 Hz) tones (50 ms duration including 5 ms rise and fall times) corresponding to a 4:1 standard to deviant ratio. The total duration of each sequence was 12 min and participants were given a short break after 6 min. Alternating short and long intervals between tones ensured that the sequences resembled typical paired stimulus SG paradigms (**Figure 1**). The intervals between the tones of a pair (intra-pair-interval, intra-PI) were 200 ms in the fully predictable isochronous sequence and between 100-300 ms in the random sequence. The intervals between pairs (inter-pair-interval, inter-PI) were 700 ms in the isochronous sequence and between 350-1050 ms in the random sequence. The random sequence was designed so that participants still perceived the tones in pairs, but the intra-PIs (100ms, 150ms, 200 ms, 250 ms, 300 ms) and inter-PIs (350 ms, 525 ms, 700 ms, 875 ms, 1050 ms) varied. The order of these time intervals was randomized using a Williams design (Williams, 1949).

**Figure 1.**
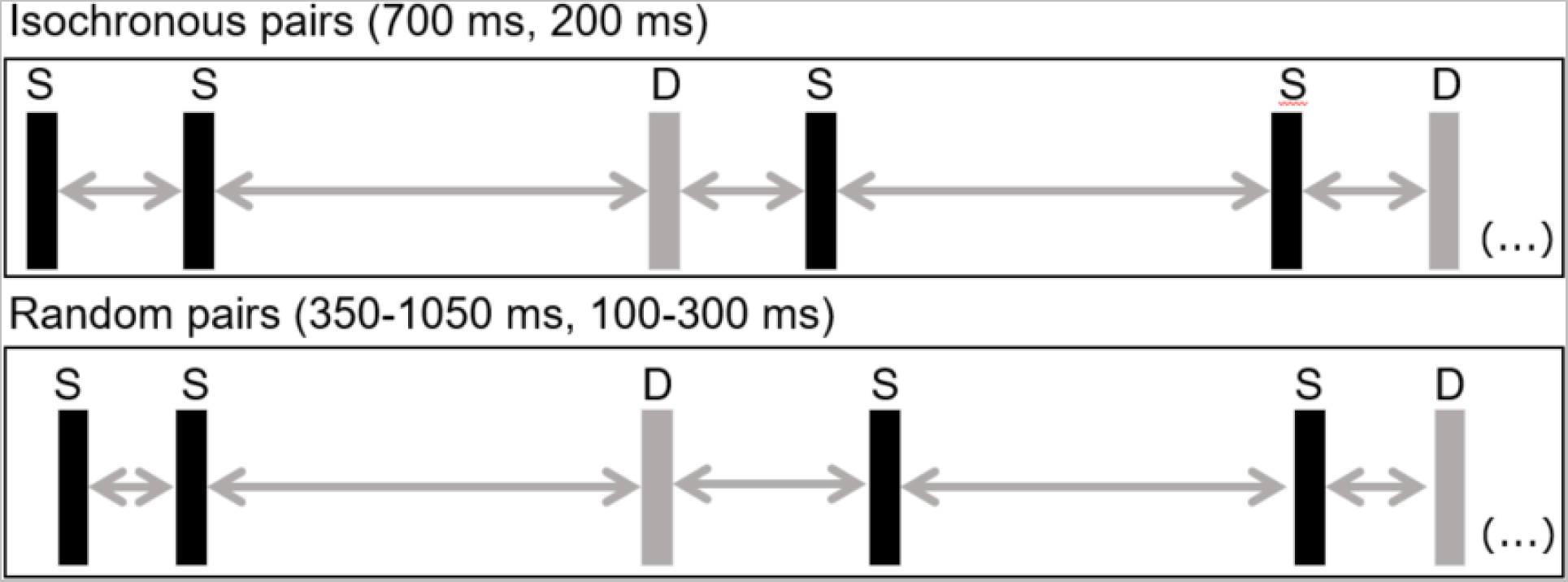
Schematic representation of the stimulus sequences. Pairs of sinusoidal tones were continuously presented in two sequences, one in which the intervals between tones and pairs were fixed (isochronous pairs) and one in which both intervals varied in duration with the constraint that shorter and longer intervals alternated (random pairs). Black bars represent standard tones (S), gray ones deviant tones (D). Standard and deviant tones differed in pitch and were presented with an overall 4:1 ratio.

### 2.4 EEG recording and pre-processing

EEG was recorded from 128 active electrodes (actiCAP, Brain Products GmbH), mounted into an elastic cap at 1000 Hz sampling rate, while impedances were kept at ≤10 kOhm. FCz was used as the online reference and the audio signal was recorded with the EEG. Data were then downsampled to 500 Hz and a bandpass filter (1-44 Hz) was applied using EEGlab (Delorme & Makeig, 2004). To detect and reject bad channels *clean_rawdata* was used (Mullen et al., 2015). Rejected channels (*M* = 5.5, *SD* = 4.9) were spherically interpolated, then the online reference was added back to the data and data were re-referenced to the average (following Foti, Hajcak, and Dien (2009)). Artifact subspace reconstruction (ASR) was performed using *clean_rawdata* and data were re-referenced again as suggested by Makoto (2022) to reorganize the data to be zero-sum across channels. Then ICA (runica; 30 pca components) was performed (Makeig, Jung, Bell, Ghahremani, & Sejnowski, 1997). IC components (*M* = 4.3, *SD* = 1.2) reflecting eye horizontal and vertical movements, muscle activity, heart rate, line noise or channel noise were rejected using *IClabel* (Pion-Tonachini, Kreutz-Delgado, & Makeig, 2019).

### 2.5 ERP analysis

Epochs lasting from −50 ms to 145 ms relative to stimulus onset were created, baseline corrected (−50 to 0 ms), and then averaged per participant. To avoid bias or double dipping about time and spatial distribution of the ERP components (Kriegeskorte, Simmons, Bellgowan, & Baker, 2009; Luck & Gaspelin, 2017), analyses followed a data-driven approach. A two-step temporal-spatial PCA (tsPCA) was performed using the EP toolkit (version 2.95) (Dien, 2010; Dien, 2012). This procedure decomposes the data based on covariances between voltages at sampling points and sampling sites and aims to identify and disentangle components that are transparently and objectively extracted (Dien & Frishkoff, 2005). Following the guidelines formulated by Dien (2012), first, a temporal PCA was performed on the averaged data, using participants, stimulus types, and recording sites as observations. A (oblique) Promax rotation was used (Hendrickson & White, 1964) and seven factors were extracted after inspection of the scree plot (Cattell, 1966) with the help of a parallel test that compares the scree plot obtained with the experimental data with one that is derived from random data (Horn, 1965). Second, a spatial (orthogonal) Infomax ICA was performed on each temporal component that survived the first step, and seven spatial components were extracted (Bell & Sejnowski, 1995; Delorme & Makeig, 2004).

### 2.6 Statistical analysis

For the demographics, independent t-tests were performed. A chi-square test was performed for sex differences and the robust counterpart of the t-test as implemented in the *WRS2* (version 1.1-4) package (Mair & Wilcox, 2020) was used when assumptions were violated. All analyses were performed in *R* (version 4.2.0) using *Rstudio* (version 2022.07.1). For the EEG data, after inspection of the time course and the spatial distribution of the components of interest, the microvolt scaled amplitudes of the max and min peaks were analyzed for two time windows (i.e., TF1SF1: 90 – 130 ms, TF3SF1: 30 – 70 ms; with TF denoting temporal factor and SF spatial factor). Subsequently, 2 (without Tinnitus vs. Tinnitus) x 2 (Isochronous vs. Random) x 2 (Standard vs. Deviant) x 2 (Position 1 vs Position 2) mixed ANOVAs were performed, separately for TF1SF1 and TF3SF1 using the *rstatix* (version 0.7.0) package (Kassambara, 2021). When checking the assumptions, some outliers were identified and two extreme outliers excluded (i.e., exceeding the interquartile range by a threefold), as they were outliers for several combinations of factors. Two participants without tinnitus were correspondingly excluded. Levene’s tests were not significant and normality was assumed based on the central limit theorem. All effects are reported as significant at *p* < .05. Effect sizes are reported as generalized eta squared (*η^2^_G_*). Non-normally distributed variables such as the Hearing loss (HL) and duration of tinnitus in months underwent square root transformation (see supplementary material for further correlation analyses).

## 3. Results

### 3.1. Demographics

There was no significant age (*t* (50) = −1.36, *p* = .18, CI [−14.25, 2.74]) or sex difference between the two groups (*χ*2 (1, *N* = 2) = 0, *p* = 1). To assess handedness, the robust t-test that is based on a two-sample trimmed mean test was performed (Yuen, 1974). The result likewise indicated no significant difference between the two groups (*t_y_* (15.45) = 0.76, *p* = .46, CI [−7.13, 3.39]). There was also no significant difference in terms of education *t* (50) = 1.68, *p* = .1 CI [−0.15, −1.67]). However, in line with previous studies, hearing loss differed between the two groups (*t* (50) = −3.26, *p* = .002, CI [−18.44, −4.4]), indicating increased hearing loss in the tinnitus group.

### 3.2 tsPCA results and selection of ERP components

The temporal-spatial PCA revealed seven temporal factors, explaining 95% of the total variance and seven spatial factors that explained 88% of the total variance. An overview of all 23 components that explained at least .5% of the total and .5% of the unique variance can be found in **Table 2**. Two temporal factors that reflected P50- or N100-like responses were selected (**Figure 2**). As the factor combinations are microvolt scaled reconstructed ERP components, P50 and N100 components of interest are referred to as P50- and N100-like responses. The first factor (TF1SF1) displayed a negative peak at channel FFC1h and explained 15.97% of unique variance. The frontocentral distribution included 42 channels (i.e., AFp1, AFp2, AF3, AFz, AF4, AFF5h, AFF1h, AFF2h, AFF6h, F5, F3, F1, Fz, F2, F4, F6, FFT7h, FFC5h, FFC3h, FFC1h, FFC2h, FFC4h, FFC6h, FFC8h, FC5, FC3, FC1, FCz, FC2, FC4, FC6, FCC5h, FCC3h, FCC1h, FCC2h, FCC4h, FCC6h, C3, C1, Cz, C2, C4), with an absolute factor loading threshold of 0.6 (Dien, 2010). This corresponds to the N100 ERP component, considering previous literature on the temporal-spatial characteristics of the N100 and visual inspection (**Figure 3**) (Davis, Mast, Yoshie, & Zerlin, 1966; Hillyard, Hink, Schwent, & Picton, 1973; Luck, 2014). Another component, TF3SF1, explained 2.07% of unique variance and its frontocentral distribution, with absolute factor loadings of 0.6, encompassed 34 electrode sites (i.e., AF3, AFz, AF4, AFF1h, AFF2h, F3, F1, Fz, F2, F4, F6, FFC5h, FFC3h, FFC1h, FFC2h, FFC4h, FFC6h, FC5, FC3, FC1, FCz, FC2, FC4, FCC5h, FCC3h, FCC1h, FCC2h, FCC4h, C3, C1, Cz, C2, CCP3h, CCP1h). Previous literature shows that the P50 has a frontocentral maximum and peaks around 40 – 80 ms (Patterson et al., 2008). Although the absolute voltage of the peak channel was negative, indicating heterogeneities across conditions, visual inspection, time course, and spatial distribution suggest that TF3SF1 corresponds to the P50 ERP component (**Figure 4**).

**Figure 2.**
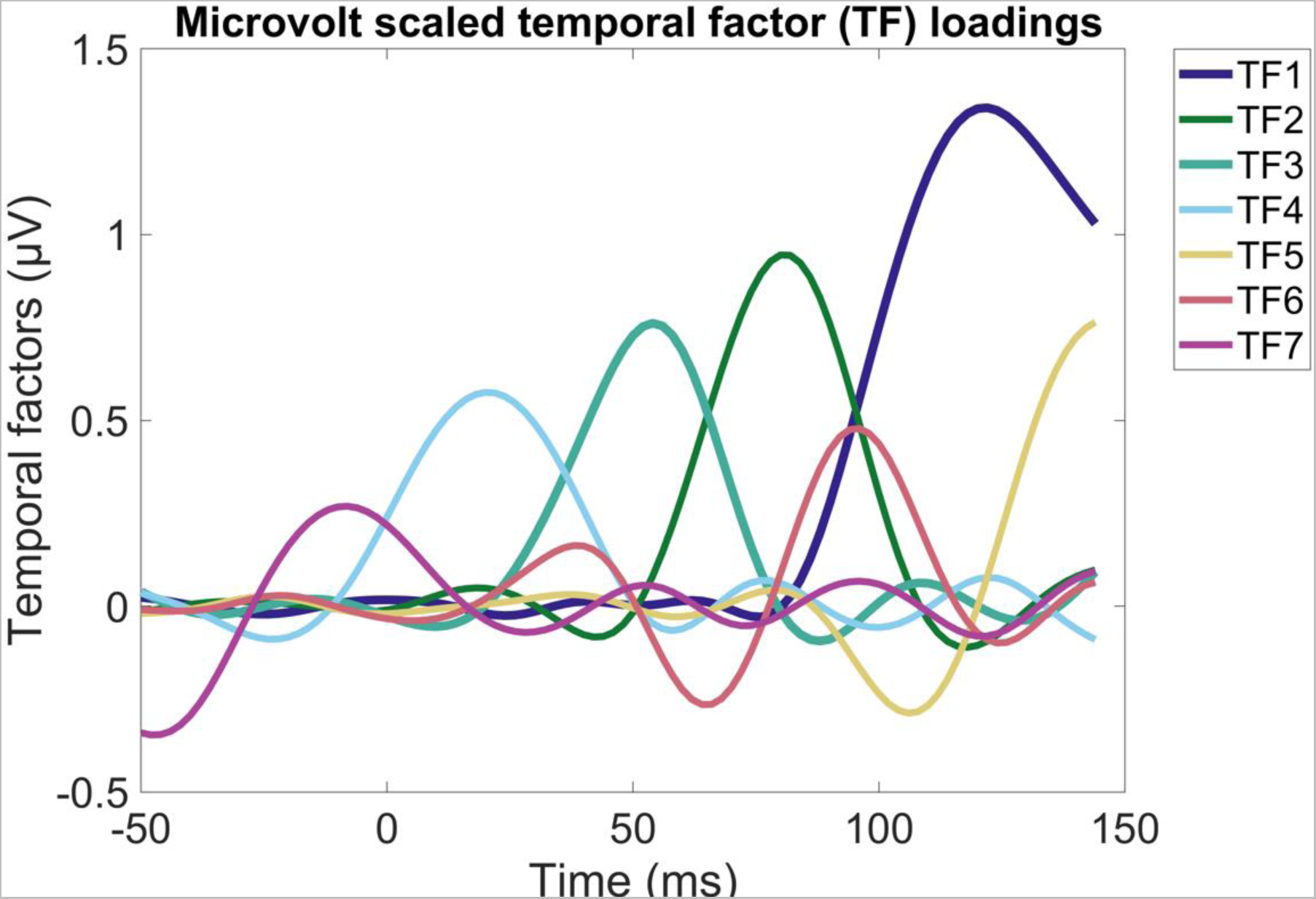
Microvolt scaled temporal factors based on temporal-spatial principal component analysis of ERP data.

**Figure 3.**
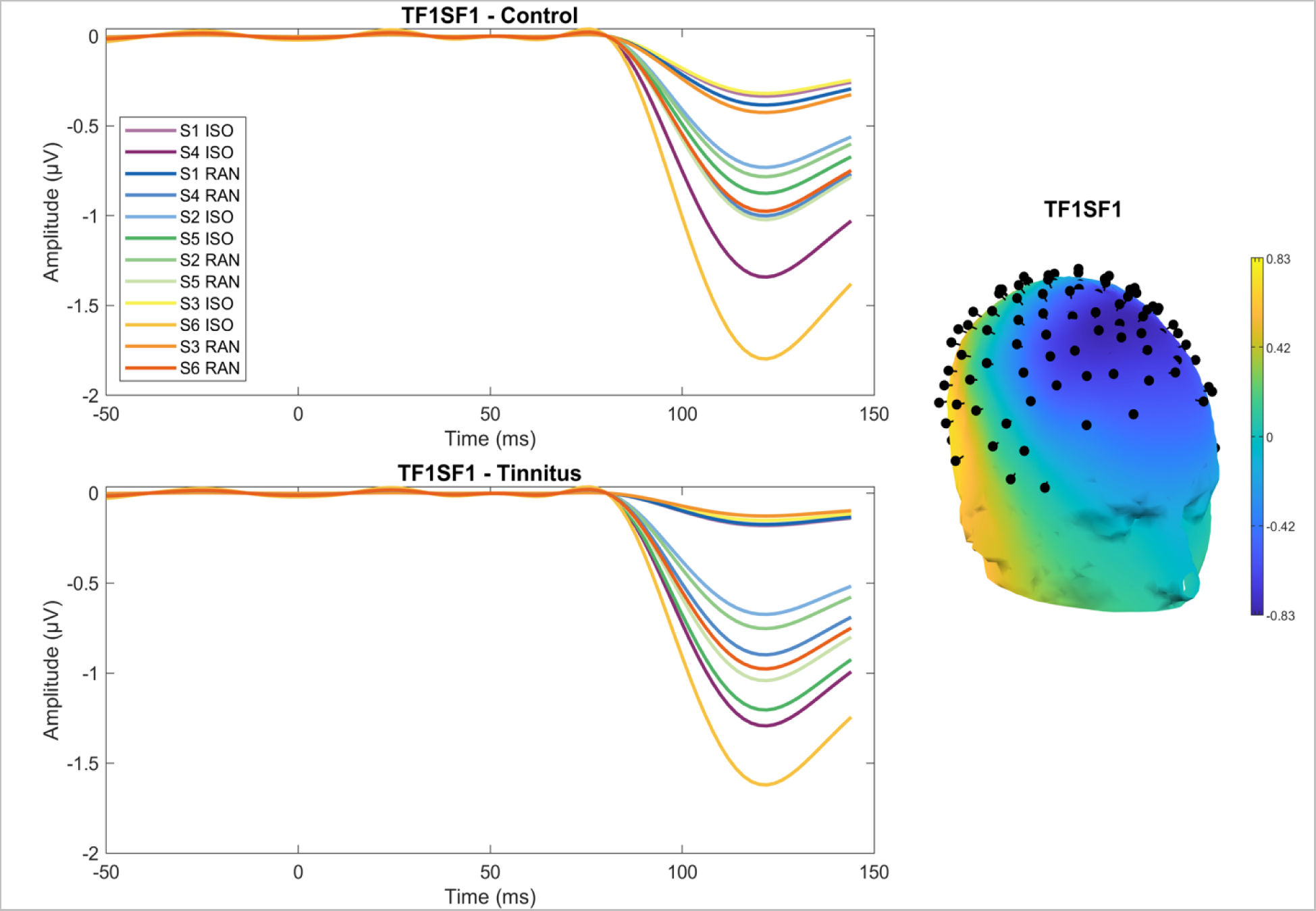
Left: Temporal-spatial factor reflecting N100-like activity for persons without tinnitus (Control, top) and persons with tinnitus (Tinnitus, bottom). Right: Topographical depiction of TF1SF1 for all conditions and stimuli, and both groups combined.

**Figure 4.**
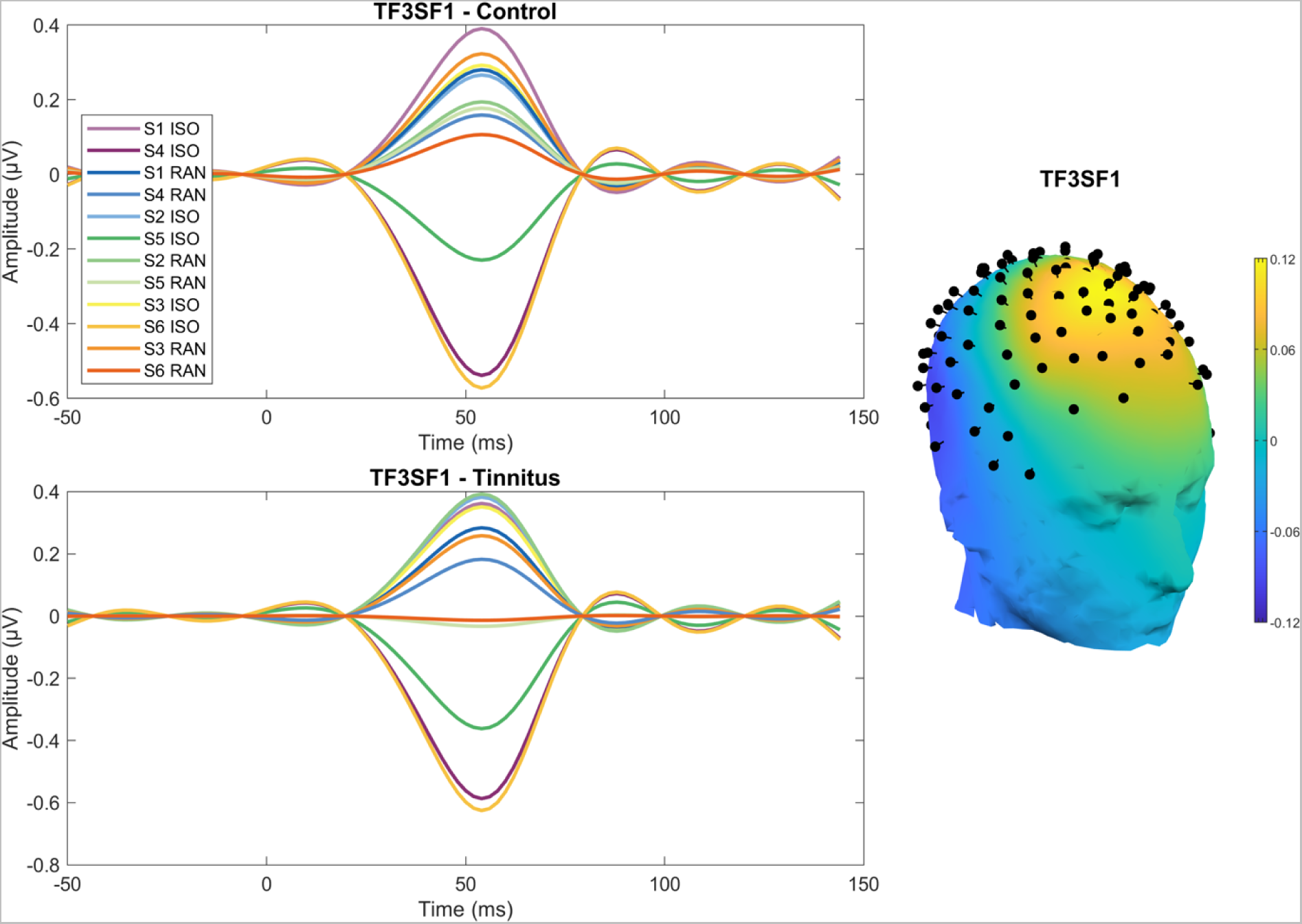
Left: Temporal-spatial factor reflecting P50-like activity for persons without tinnitus (Control, top) and with tinnitus (Tinnitus, bottom). Right: Topographical depiction of TF3SF1 for all conditions and stimuli, for both groups combined.

**Table 2.** Principal components explaining at least .5% of the total variance and .5% of the unique variance.

| Principal component | Peak latency (ms)<sup>a</sup> | Peak channel (-)<sup>b</sup> | Peak channel (+)<sup>b</sup> | Peak<sup>c</sup> | Total variance explained | Unique variance explained |
| --- | --- | --- | --- | --- | --- | --- |
| TF1SF1 | 122 | FFC1h | O10 | - | 26.53% | 15.97% |
| TF1SF2 | 122 | CCP1h | Fp1 | + | 6.97% | 4.11% |
| TF1SF3 | 122 | Fp1 | TP9 | + | 5.19% | 3.09% |
| TF1SF4 | 122 | FTT9h | FTT8h | + | 2.39% | 1.44% |
| TF1SF5 | 122 | O10 | CCP5h | + | 1.09% | 0.67% |
| TF1SF6 | 122 | FC5 | PO7 | + | 0.97% | 0.60% |
| TF2SF1 | 80 | FFC1h | O10 | - | 7.63% | 3.78% |
| TF2SF2 | 80 | CP2 | Fp1 | + | 2.28% | 1.09% |
| TF2SF3 | 80 | Fp1 | TP9 | + | 1.67% | 0.86% |
| TF2SF4 | 80 | P7 | FTT8h | + | 1.41% | 0.71% |
| TF2SF5 | 80 | PO9 | CCP5h | + | 1.17% | 0.53% |
| TF2SF6 | 80 | FFC5h | PPO9h | - | 1.11% | 0.52% |
| TF3SF1 | 54 | FFC1h | PPO10h | - | 4.62% | 2.07% |
| TF3SF2 | 54 | F9 | Pz | - | 1.57% | 0.68% |
| TF3SF3 | 54 | O9 | Fp1 | + | 1.44% | 0.63% |
| TF3SF4 | 54 | P7 | C6 | - | 1.22% | 0.55% |
| TF4SF1 | 20 | FFC1h | PO8 | + | 2.69% | 1.51% |
| TF4SF2 | 20 | F10 | CPP1h | + | 1.43% | 0.80% |
| TF4SF3 | 20 | TP7 | C4 | - | 0.98% | 0.55% |
| TF5SF1 | 144 | TPP9h | FFC1h | + | 3.32% | 2.58% |
| TF5SF2 | 144 | F10 | Pz | + | 0.84% | 0.66% |
| TF6SF1 | 96 | FFC3h | TPP9h | + | 2.39% | 1.29% |
| TF7SF1 | -48 | PPO5h | AFF2h | - | 0.80% | 0.64% |

<sup>a</sup> Time point with the largest voltage.
<sup>b</sup> Channel with the greatest voltage for positive and negative voltages.
<sup>c</sup> Whether the negative or positive peak channel had a greater absolute voltage.

#### 3.2.1 P50-like activity (TF3SF1)

There were significant main effects of *temporal structure* (i.e., smaller values for the isochronous condition), *F*(1,48) = 20.54, *p* < .0001, *η^2^_G_* = .064, *formal structure* (i.e., larger values for deviants), *F*(1,48) = 49.02, *p* < .0001, *η^2^_G_* = .087, and *position* (i.e., smaller values for position 2), *F*(1,48) = 150.25, *p* < .0001, *η^2^_G_* = .337. Additionally, there were interaction effects of *temporal structure* and *formal structure*, *F*(1,48) = 4.64, *p* = .036, *η^2^_G_* = .005, and of *temporal structure* and *position*, *F*(1,48) = 38.52, *p* < .0001, *η^2^_G_* = .085. Post-hoc analyses consisted of Bonferroni corrected simple pairwise comparisons and confirmed significant differences between the isochronous and random condition for deviant, t(49) = −4.07, *p* < .001, and standard, t(49) = −4.43, *p* < .0001, tones. Deviant and standard responses also differed within the isochronous t(49) = 4.88, *p* < .001 and the random t(49) = 6.42, *p* < .001 condition. (**Figure 5**). For the *temporal structure* x *position interaction*, post-hoc tests showed significant differences between the isochronous and random condition for position 2, t(49) = −6.01, *p* < .001, but not for position 1, t(49) = 0.821, *p* = .416 responses. In addition, significant differences between position 1 and position 2 were obtained for the isochronous, t(49) = 11.2, *p* < .001 and the random, t(49) = 7.2, *p* < .001 condition (**Figure 5**).

**Figure 5.**
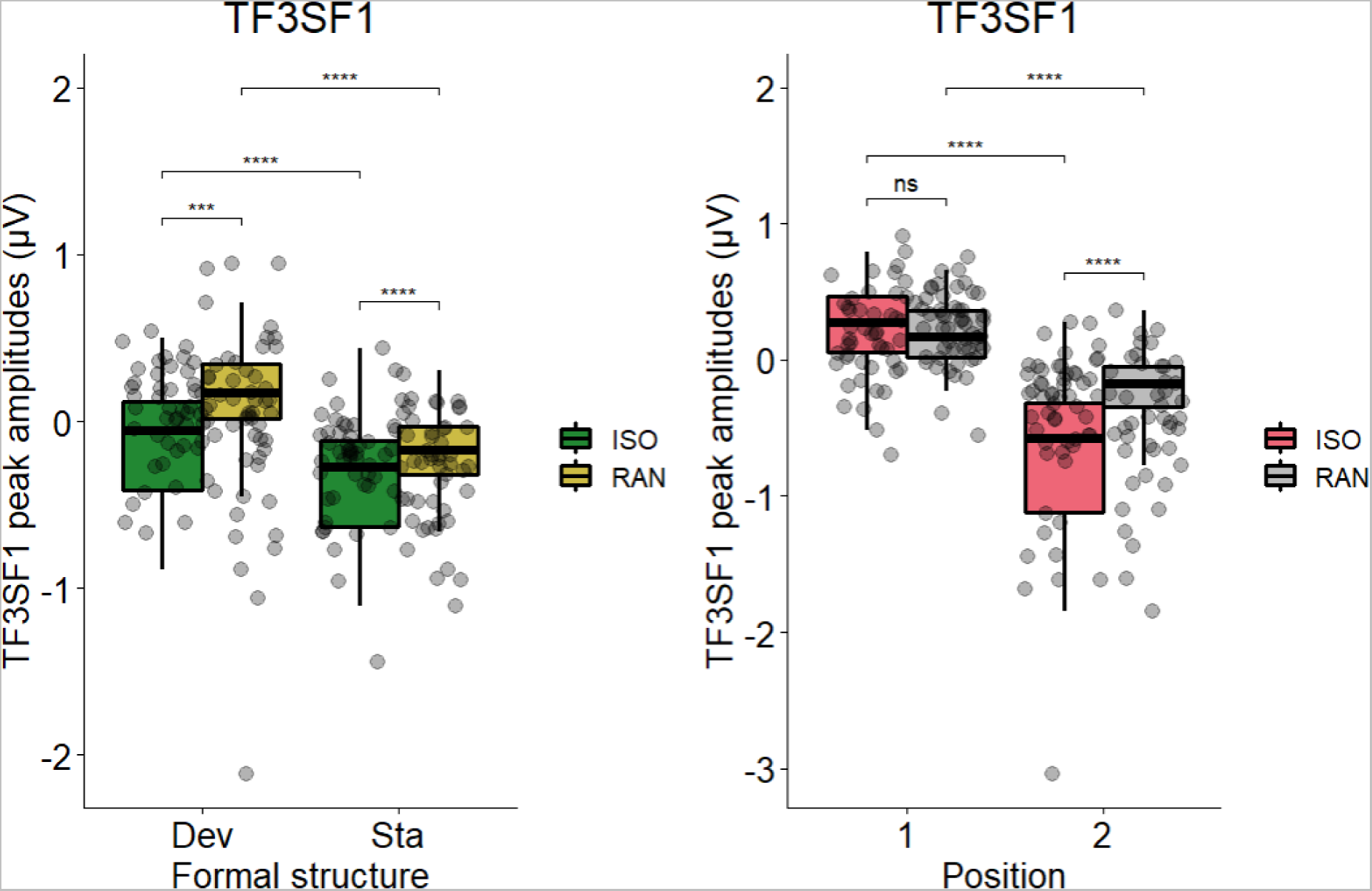
Peak amplitudes in microvolts for the third temporal and first spatial factor (TF3SF1) for the *temporal structure* x *formal structure* (left) and *temporal structure* x *position* (right) interaction. Abbreviations: Sta: standard, Dev: deviant, ISO: isochronous condition, RAN: random condition, *** *p* ≤ .001.

#### 3.2.2 N100-like activity (TF1SF1)

As expected, there were significant main effects of *temporal structure* (i.e., isochronous versus random) indicating more negative values for the isochronous condition, *F*(1,48) = 21.96, *p* < .0001, *η^2^_G_* = .028, *formal structure* (i.e., standards versus deviants), reflecting more negative values for deviants, *F*(1,48) = 45.12, *p* < .0001, *η^2^_G_* = .057, and *position* (i.e., position 1 versus position 2), indicating more negative values for position 2, *F*(1,48) = 91.46, *p* < .0001, *η^2^_G_* = .248. Additionally, there were interactions of *temporal structure* and *formal structure*, *F*(1,48) = 19.42, *p* < .0001, *η^2^_G_* = .01, *temporal structure* and *position*, *F*(1,48) = 33.34, *p* < .0001, *η^2^_G_* = .038, and *formal structure* and *position*, *F*(1,48) = 11.17, *p* = .002, *η^2^_G_* = .008. For the *temporal structure* x *position interaction*, post-hoc tests showed significant differences between the isochronous and random condition for position 2, t(49) = −7.38, *p* < .001, but not for position 1, t(49) = 0.585, *p* = .561 responses. In addition, significant differences between position 1 and position 2 were obtained for the isochronous, t(49) = 9.96, *p* < .001 and the random, t(49) = 6.54, *p* < .001 condition (**Figure 6**). Finally, three-way interactions of *temporal structure* x *formal structure* x *position* was significant, *F*(1,48) = 33.26, *p* < .0001, *η^2^_G_* = .013, and *group* x *temporal structure* x *formal structure interaction* were significant, *F*(1,48) = 4.65, *p* = .036, *η^2^_G_* = .002. Post-hoc analyses consisted of a Bonferroni corrected simple-two way interaction at each level of *group*, resulting in significant main effects of *temporal structure*, *F*(1,22) = 9.94, *p* = .03, *η^2^_G_* = .049, and *formal structure*, *F*(1,22) = 19.2, *p* < .01, *η^2^_G_* = .072, in persons without tinnitus, as well as in persons with tinnitus (i.e., *temporal structure*: *F*(1,26) = 12.1, *p* = .012, *η^2^_G_* = .031, *formal structure*: *F*(1,26) = 27.1, *p* < .001, *η^2^_G_* = .087). However, only persons without, *F*(1,22) = 13.1, *p* = .012, *η^2^_G_* = .033, but not with tinnitus, *F*(1,26) = 4.91, *p* = .216, *η^2^_G_* = .003, showed the *temporal structure* x *formal structure interaction*, that remained significant after Bonferroni correction. For persons without tinnitus, a simple simple main effect further indicated a significant difference between temporal conditions selectively for the deviant tones, *F*(1,22) = 14.9, *p* < .001, *η^2^_G_* = .132, but not for standard tones, *F*(1,22) = 0.41, *p* = .53, *η^2^_G_* = .002. Finally, following up on the effect of temporal structure for deviant tones in persons without tinnitus, Bonferroni corrected simple simple pairwise comparisons showed a significant difference between the isochronous and the random condition for deviant tones for persons without tinnitus, t(22) = −3.86, p < .001 (**Figure 7**).

**Figure 6.**
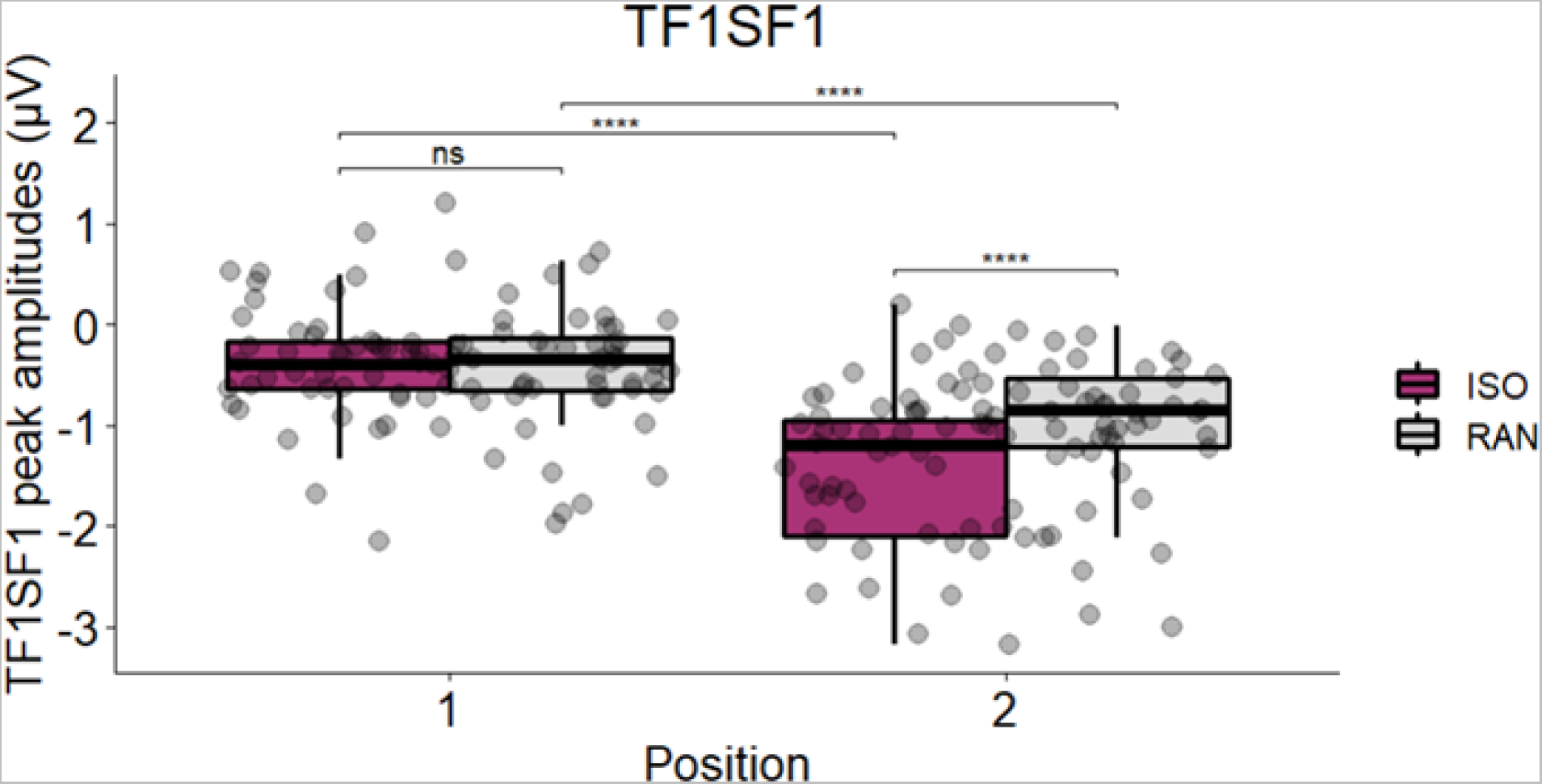
Peak amplitudes in microvolts for the first temporal and first spatial factor (TF1SF1) *temporal structure* x *position* interaction. Abbreviations: Sta: standard, Dev: deviant, ISO: isochronous condition, RAN: random condition, *** *p* ≤ .001.

**Figure 7.**
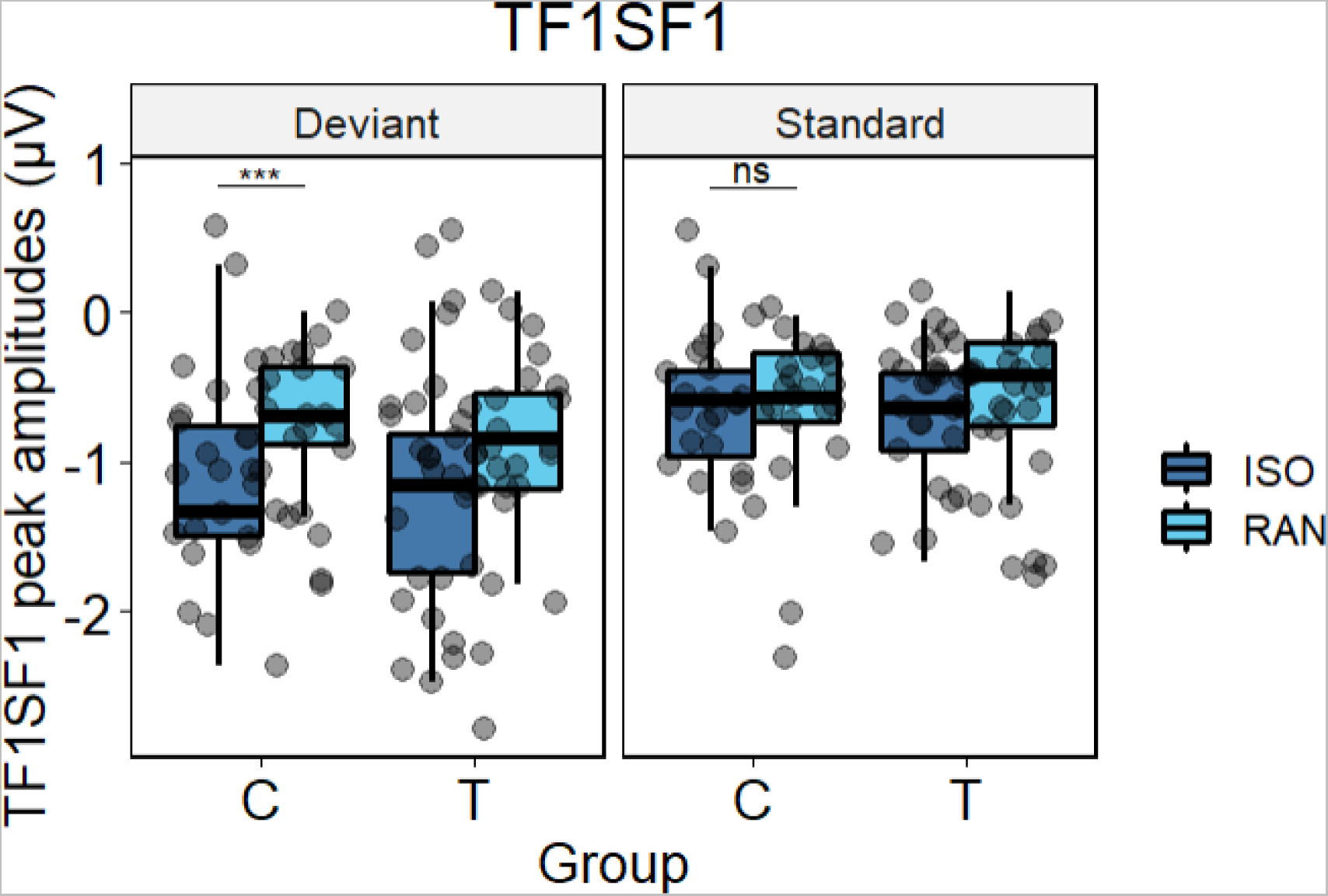
Peak amplitude differences in microvolt for the first temporal and spatial factor (TF1SF1) for the simple pairwise comparisons. Abbreviations: ISO: isochronous condition, RAN: random condition, C: control Group/persons without tinnitus, T: persons with tinnitus, *** *p* ≤ .001.

## 4. Discussion

This study investigated modulations of the P50- and N100-like activity in response to three different dimensions of stimulus predictability in persons with and without tinnitus. Manipulations of predictability altered the formal and temporal structure as well as the position of tones presented in pairs. This binary grouping made the setup comparable to classical binary SG paradigms. P50-like activity indicated expected effects of formal structure, temporal structure, and position in both groups. In other words, P50 amplitudes were smaller for standard than deviant tones, smaller for isochronous than random stimulus timing, and smaller in response to the second than the first tone of tone pairs, confirming a classical SG effect in both groups. These results confirm the effectiveness of a paradigm to simultaneously assess formal-, temporal- and position predictability. Additionally, there was a significant difference between the temporal conditions for the second tone in pairs, while this was not the case for the first tone. This again indicates a stronger amplitude reduction in the isochronous than the random condition in both groups. This overall pattern repeated in the N100-like activity, while globally, amplitudes for the isochronous sequence were more negative than for the random sequence. However, amplitudes in N100-like activity were larger for deviant than for standard tones in the isochronous timing condition in persons without tinnitus only. Previous research reported that participants respond faster when stimuli are presented in a temporally regular than irregular context (Lange, 2009; Rohenkohl, Cravo, Wyart, & Nobre, 2012) and that temporal predictability facilitates stimulus detection (Lawrance, Harper, Cooke, & Schnupp, 2014). At the neurophysiological level, the suppression of early ERP responses to expected stimuli is a well-established phenomenon (Bendixen et al., 2012; Costa-Faidella, Baldeweg, Grimm, & Escera, 2011; Lange, 2009; Schwartze et al., 2013). Therefore, the current findings for temporal predictability are in line with these previous findings.

However, the current study did not reveal differences for position predictions (SG) between persons with and without tinnitus in P50- and N100-like activity. Previously, Campbell et al. (2018) found a correlation for tinnitus severity, suggesting decreased gating in the Pa component when tinnitus burden increased. However, similar to the current results, they reported no group differences in SG for the Pa, P50, N100 or P200 components (Campbell et al., 2018). Similar P50 results were reported by Dornhoffer et al. (2006). Compared to the current study, participants in the Campbell et al. (2018) study had similar hearing thresholds but were younger (i.e., on average between 20 and 22 years) and their tinnitus handicap was very low (i.e., 0 – 14). A tinnitus handicap inventory (THI) score can range between 0 - 100 and scores between 0 - 16 indicate no or only a slight handicap (Lee, Ra, & Kim, 2014; Newman et al., 1996). Here, we administered the TQ and not the THI, even though both instruments have shown high convergent validity (Baguley, Humphriss, & Hodgson, 2000). In addition, the paradigms differed in terms of the intra-PI (500 ms) and the inter-PI (7 s) (Campbell et al., 2018). As the current paradigm used shorter intra-PIs and inter-PIs and to avoid double dipping, we applied tsPCA, a data-driven analysis that can delineate overlapping processes and can therefore enhance the signal-to-noise ratio (Dien, 2012; Foti et al., 2009). Moreover, in the current study, N100-like activity was frontally located, which might be explained by age of in the current participant sample (Paitel & Nielson, 2021). In Dornhoffer et al. (2006), the age range was similar to the current study. However, another tinnitus severity questionnaire (i.e., tinnitus severity index questionnaire) was administered. With different intra-PIs (i.e., 250 ms, 500 ms, 1000 ms), there were no group differences for P50 SG (Dornhoffer et al., 2006). Overall, the current study thus produced similar results as Campbell et al. (2018) and Dornhoffer et al. (2006), despite methodological differences between the studies. This confirms the robustness of the results, suggesting that SG dysfunctions in tinnitus are likely more subtle than assumed.

Most importantly, we observed SG effects in persons with and without tinnitus, indicating successful processing of position predictions and an associated reduced response. Early SG research focused on persons with schizophrenia, showing that SG is dysfunctional in this group (Adler et al., 1982; Patterson et al., 2008). Other research observed a similar pattern in persons with Alzheimer’s disease (Jessen et al., 2001) or ADHD (Davies, Chang, & Gavin, 2009; Holstein et al., 2013). Similar to the current and Campbell et al. (2018) findings, unaltered SG in persons with tinnitus was also reported in high-functioning children along the autism spectrum (Kemner, Oranje, Verbaten, & Engeland, 2002; Orekhova et al., 2008), and in patients with obsessive-compulsive disorder (de Leeuw, Oranje, van Megen, Kemner, & Westenberg, 2010). In the current study, we now show that for the P50-like and the N100-like activity, predictions regarding formal structure and position are intact in both groups in a temporally predictable context, indicating intact binary auditory stimuli filter mechanisms in persons with tinnitus.

For the N100-like activity, however, a different pattern emerged for deviance processing in the two timing conditions in persons with and without tinnitus. This observation can be linked to different attentional processing for deviant events in persons with and without tinnitus. Altered attention in persons with tinnitus has previously been proposed by Roberts et al. (2013). Roberts et al. (2013) proposed a qualitative model, in which attention allocation changes following a mismatch between the incoming auditory input and the sound representation generated in the auditory cortex. In addition, a meta-analysis investigated behavioral and electrophysiological measures of attention in persons with tinnitus and confirmed that later attentional processes are altered in persons with tinnitus as indicated by reduced mismatch negativity (MMN) and P300 amplitudes, although the heterogeneity of the tinnitus populations and methods limit precise interpretations of the underlying mechanisms (Vasudevan, Ganapathy, Palaniswamy, Searchfield, & Rajashekhar, 2021). When assessing the N100 component in persons with tinnitus, reduced N100 amplitudes were observed for persons with distressing (i.e., decompensated) tinnitus (Jacobson & McCaslin, 2003). Moreover, persons with low and high tinnitus-related distress showed differences in N100 activity (Delb et al., 2008). More specifically, in an unattended condition, in which participants had to ignore tones and think of something pleasant, differences were found between persons with low tinnitus distress and high tinnitus distress (more negative N100 for high distress) and between persons without tinnitus and high tinnitus distress (more negative N100 for high distress). In the attended condition, N100 amplitudes of persons with high distress were more negative than without tinnitus (Delb et al., 2008). Other research on auditory attention in tinnitus (Roberts et al., 2013), suggests facilitatory cholinergic neuromodulation in cortico-subcortical pathways at the level of the ventromedial prefrontal cortex and basal forebrain, which could reinforce aberrant neural synchrony in persons with tinnitus. Interestingly, it was reported that activity in fronto-parietal regions differs in persons with tinnitus when they are presented with tinnitus-specific frequency sounds as opposed to a control frequency (Salvari et al., 2023). These findings may indicate why a more efficient adaptation to deviant tones was observed in persons without tinnitus than in persons with tinnitus and suggests altered auditory attention allocation in response to deviant tones in the tinnitus group. Along these lines, the current findings may indicate a shifting of attentional bias toward the tinnitus percept or altered redirection of selective auditory attention away from it. Future research should therefore delineate how auditory predictions are influenced by tinnitus-specific frequencies (i.e., regularly 2-4 kHz in tinnitus linked to noise induced hearing loss (Eggermont & Roberts, 2004)) for the P50- and N100-like components, respectively.

Lastly, when working with a clinical population such as persons with tinnitus, controlling for hearing loss, age, and/or hyperacusis at the same time is challenging. Tinnitus is more prevalent in older persons and often accompanied by some degree of hearing loss (Axelsson & Ringdahl, 1989; Knipper, Van Dijk, Nunes, Rüttiger, & Zimmermann, 2013; Nelson & Chen, 2004), although it can also occur without altered hearing thresholds (Roberts et al., 2010; Weisz, Hartmann, Dohrmann, Schlee, & Norena, 2006). The average hearing loss in the current tinnitus sample was very mild and associated with no impairment at all or with slight difficulties (Olusanya, Davis, & Hoffman, 2019). Research that includes persons with tinnitus with and without hearing loss and comparisons to hearing-matched persons without tinnitus remains scarce. One EEG study that investigated prediction errors for auditory false perceptions and phantom perceptions in persons with tinnitus assigned participants to groups with mild or severe hearing loss and compared them to a group of persons with schizophrenia (Ahn et al., 2022). The results showed that auditory ERP responses for self-generated sounds were not suppressed in persons with tinnitus + severe hearing loss (n = 15) and persons with schizophrenia (n = 10), whereas the typical suppression was observed in persons without tinnitus (n = 23) and in persons with tinnitus + mild hearing loss (n = 8) (Ahn et al., 2022). However, the degree of hearing loss in the tinnitus + hearing loss group exceeded the level of hearing loss in the current group, while also no full high-frequency audiometry was performed here. Similarly, an MRI study that compared persons with hearing loss, tinnitus + hearing loss, and controls showed that tonotopic maps for the hearing loss + tinnitus were more similar to controls relative to persons with hearing loss only (Koops, Renken, Lanting, & van Dijk, 2020). Therefore, it was concluded that tinnitus might be a side product of dampened cortical reorganization and not the result of it (Koops et al., 2020). However, considering the overall mild hearing loss in the current sample, it is likely that hearing loss only had a minor influence on the results reported here.

The results show that the adopted paradigm permits assessing three dimensions of auditory prediction (i.e., formal-, temporal-, and position). Classic position-based SG was not altered in persons with and without tinnitus for the P50-like activity. However, for the N100-like activity, deviance processing was further modulated by temporal regularity only in persons without tinnitus but not in persons with tinnitus. Hence, it seems likely that temporally regular and thus fully predictable stimulation facilitates deviance processing in persons without tinnitus and that this mechanism is altered in persons with tinnitus. Auditory filtering as indexed by classical SG effects for binary auditory stimuli seems thus not substantially different in persons with and without tinnitus. It rather seems that tinnitus alters attention-allocation in response to the deviant, i.e., unpredicted or at least less predictable auditory events. This finding might indicate a shifting attentional bias towards the tinnitus sound that may be accompanied by dysfunctional allocation of selective auditory attention to other sounds.

## CRediT author statement

P. B.: Formal analysis, Investigation, Writing – Original Draft.

J. V. P. D.: Investigation, Writing – Review & Editing.

J. H. M. v. d. E.: Investigation, Writing – Review & Editing.

J. V. S.: Writing – Review & Editing.

M. L. F. J.: Conceptualization, Writing – Review & Editing, Supervision.

S. A. K.: Conceptualization, Writing – Review & Editing, Supervision.

M. S.: Conceptualization, Methodology, Writing – Review & Editing, Supervision.

## Funding

PB was funded by a PhD scholarship from the German Academic Scholarship Foundation (Studienstiftung des deutschen Volkes).

## Conflict of interest disclosure

The authors declare no conflict of interest.

## Notes

### Competing Interest Statement

The authors have declared no competing interest.

