## supplementary material for "Parallel EEG assessment of different sound predictability levels in tinnitus"

1. Relationship between classical sensory gating, HL, TQ and duration of tinnitus

1.1. Methods

Pearson correlations were performed on the standard – standard tone pair in the isochronous condition as implemented in the *rstatix* package. SG was quantified in terms of peak amplitude differences (Position 1 – Position 2). In general, SG can be quantified by calculating the SG ratio, or the SG difference, while the latter is recommended (Rentzsch, Jockers-Scherübl, Boutros, & Gallinat, 2008; Smith et al., 1994).

1.2. Results

1.2.1. TF1SF1

When correlating the SG difference scores with the amount of average HL per subgroup, a non-significant positive relation between the SG values and the amount of hearing loss was observed. In the control group improved SG correlated non-significantly with increased HL (*r* (21) = .35, *p* = .1, CI [-.07, 0.66]), similarly to the tinnitus group(*r* (25) = .21, *p* = .3, CI [-.19, .55]). For the tinnitus burden, assessed by the TQ in the tinnitus group, SG values exhibited a non-significant negative relation between SG difference scores and TQ, (*r* (25) = -.2, *p* = .33, CI [-.54, .2]). When focusing on the duration of tinnitus (in months) and SG, a non-significant positive correlation was found for the tinnitus group, *r* (25) = .16, *p* = .42, CI [-.23, .51]).

1.2.2. TF3SF1

For the TF3SF1 component a similar pattern was observed regarding the directions and the level of significance of the correlations. In the control group improved SG correlated non-significantly with increased HL (*r* (21) = .36, *p* = .095, CI [-.07, 0.67]), similarly to the tinnitus group(*r* (25) = .03, *p* = .88, CI [-.35, .41]). For the tinnitus burden, assessed by the TQ in the tinnitus group, SG values exhibited a non-significant negative relation between SG difference scores and TQ, (*r* (25) = -.17, *p* = .41, CI [-.51, .23]). When focusing on the duration of tinnitus (in months) and SG, a non-significant negative correlation was found for the tinnitus group, *r* (25) = -.016, *p* = .94, CI [-.39, .37]).
